# Imputing Single-cell RNA-seq data by combining Graph Convolution and Autoencoder Neural Networks

**DOI:** 10.1101/2020.02.05.935296

**Authors:** Jiahua Rao, Xiang Zhou, Yutong Lu, Huiying Zhao, Yuedong Yang

## Abstract

Single-cell RNA sequencing technology promotes the profiling of single-cell transcriptomes at an unprecedented throughput and resolution. However, in scRNA-seq studies, only a low amount of sequenced mRNA in each cell leads to missing detection for a portion of mRNA molecules, i.e. the dropout problem. The dropout event hinders various downstream analysis, such as clustering analysis, differential expression analysis, and inference of gene-to-gene relationships. Therefore, it is necessary to develop robust and effective imputation methods for the increasing scRNA-seq data. In this study, we have developed an imputation method (GraphSCI) to impute the dropout events in scRNA-seq data based on the graph convolution networks. The method takes advantage of low-dimensional representations of similar cells and gene-gene interactions to impute the dropouts. Extensive experiments demonstrated that GraphSCI outperforms other state-of-the-art methods for imputation on both simulated and real scRNA-seq data. Meanwhile, GraphSCI is able to accurately infer gene-to-gene relationships by utilizing the imputed matrix that are concealed by dropout events in raw data.

## Background

Compared to bulk cell RNA-sequencing^1^ (RNA-seq), single cell RNA sequencing technology^2^ (scRNA-seq) has greatly promoted the profiling of transcriptomes at single cell level and helped researchers to improve understanding of complex biological questions. It allows people to study cell-to-cell variability at a much higher throughput and resolution, such as studies of cell heterogeneity, differentiation and developmental trajectories^3^.

Despite its improvements, various technical deviations occurred due to the upgrade of sequencing techniques from bulk samples to single cells. Typically, the low RNA capture rate and sequencing efficiency leads to a large proportion of expressed genes with a false zero counts in some cells, defined as ‘dropout’ event^4, 5^. Although many of the zero counts represent the true absence of gene expression in specific cells, a considerable fraction is due to the dropout phenomenon where a truly expressed gene is undetected in some cells, resulting in zero or low read counts. Therefore, it is important to note the distinction between the truly expressed zeroes and the false zeroes in statistical analysis. Not all zeroes can be considered as the missing values to be imputed.

Furthermore, most downstream analysis on scRNA-seq data, such as cell clustering analysis, differential expression (DE) analysis and cell developmental trajectories, rely on the “corrected” gene expressions in each cell. As a result, methods such as MAGIC^6^, SAVER^7^, scImpute^8^ and DCA^9^ have been developed to correct the false zero read counts in order to recover true expression levels in scRNA-seq data. These approaches estimate “corrected” gene expressions by borrowing information across similar genes or cells. For example, MAGIC^6^ imputes gene expression data for each gene across similar cells based on Markov transition matrix, while SAVER^7^ takes advantage of gene-to-gene relationships by using Bayesian approach to infer the denoised expression. Both MAGIC^6^ and SAVER^7^ would recover the expression level of each gene in each cell including those unaffected by dropout events. ScImpute^8^, on the other hand, determines the dropout entries based on a mixture model and imputes only the likely drop-out entries across similar cells. These methods mentioned above fail to learn the non-linear relationships and the count structures in the scRNA-seq data. Thus, DCA^9^ proposes an imputation method based on an Autoencoder, a kind of deep neural networks used to reconstruct data in an unsupervised manner. In the imputation of each cell, DCA^9^ minimizes the zero-inflated negative binomial (ZINB)^10^ model-based loss function to learn gene-specific distribution in scRNA-seq data.

However, these existing imputation methods for scRNA-seq aim at learning the similarity of cells or genes only but not genes and cells simultaneously, resulting in the fact that they can’t retain biological variation across cells or genes. For example, the gene is truly not expressed due to gene regulation, but imputed by similar cells, which makes it difficult to study cell-to-cell variation and downstream analysis. This means that our imputation method not only needs to take advantage of the information between similar cells but also gene-to-gene relationships.

Accordingly, in this paper, we developed a **S**ingle-**C**ell **I**mputation method that combines **Graph** convolution network and Autoencoder neural networks, called GraphSCI, to impute the drop-out events in scRNA-seq by systematically integrating the gene expression with gene-to-gene relationships. We will use gene-to-gene relationships as prior knowledge to recover gene expression in a single cell because gene-to-gene interactions are likely to affect gene expression sensitively. The gene-to-gene relationships can be regarded as a graph, in which the gene is the node and the edge is the relationship. As a consequence, the imputation task of gene expression can be converted into the node recovering problem on graphs.

Graph convolution network^11^ (GCN) is a very powerful neural network architecture for machine learning on graphs. It was designed to learn hidden layer representations that encode both local graph structure and features of nodes and edges. A number of recent studies describe applications of graph convolution network such as node recovering problem^12-15^. Inspired by the co-embedding attributed network^12^, we combine graph convolution network and auto-encoder neural network to systematically learn the low-dimensional embedded representations of genes and cells. Graph convolution network exploits the spatial feature of gene-to-gene relationships effectively while Autoencoder neural network learns the non-linear relationships of cells and count structures of scRNA-seq data. And the deep learning framework finally reconstructs gene expressions by integrating gene expressions and gene-to-gene relationships dynamically in the backward propagation of neural networks. Fig. 1 is the architecture of our GraphSCI model.

**Figure 1.**
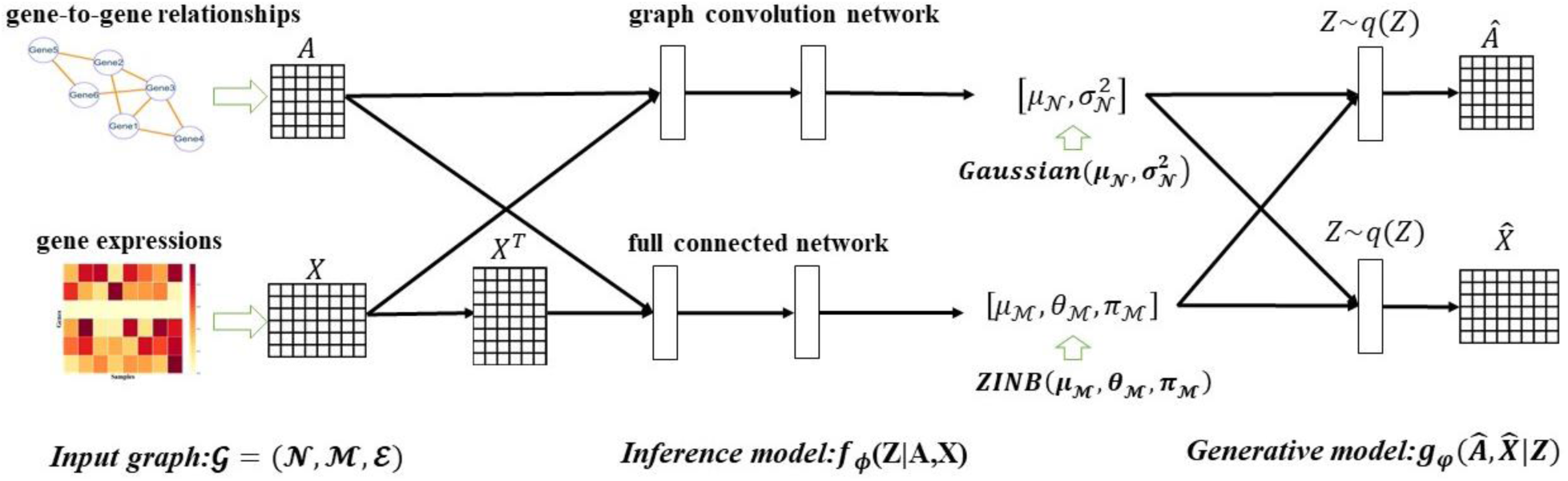
The architecture of GraphSCI model. The input of GraphSCI is the gene expression matrix and the gene-to-gene relationships. The Inference model ***f***_***ϕ***_ is to learn the low-dimensional representations of genes and cells based on a combination of graph convolution network and Autoencoder neural network. The Generative model ***g***_***φ***_ utilizes the posterior distributions to reconstruct gene expression and gene-to-gene relationships respectively.

## Results

### GraphSCI identifies cell types in simulated data

In order to assess our method, we followed the same way as the previous study^9^ to construct two simulated datasets by Splatter^16^ package: (1) 2000 cells belonging to two types clustered by expression data of 3000 genes (namely SIM-T2) and (2) 3000 cells belonging to six types of cells clustered by expression data of 5000 genes (namely SIM-T6). On the SIM-T2 data with a simpler case, GraphSCI achieved imputed expression with a mean absolute error of 0.226, which is 21.24% lower than 0.274 by DCA. We further used the imputed gene expression to cluster the cells by K-Means algorithm. As shown in Fig 2A, GraphSCI achieved 0.977, 0.994, 0.609 for ARI, CA, and SC values with standard deviation of 0.0038, 0.0043, 0.0024, respectively. These results are significantly better than 0.914, 0.925 and 0.582 achieved by DCA with P-value of 0.2292. By comparison, the clustering over original expression data without any imputation achieved 0.716, 0.508, 0.342 for ARI, CA and SC, respectively with P-value of 0.0034. The P-values indicated that although GraphSCI was not significantly better than DCA, it is significantly better than the original data. Hence it is fair to conclude that both GraphSCI and DCA are competitive on the SIM-T2 dataset. Fig 2B shows 2000 cells by using the first two principle components obtained from PCA^17^. Obviously, GraphSCI clearly separate two types of cells, while DCA has a small number of cells mixed together. The original data can’t separate the cells at all.

**Figure 2.**
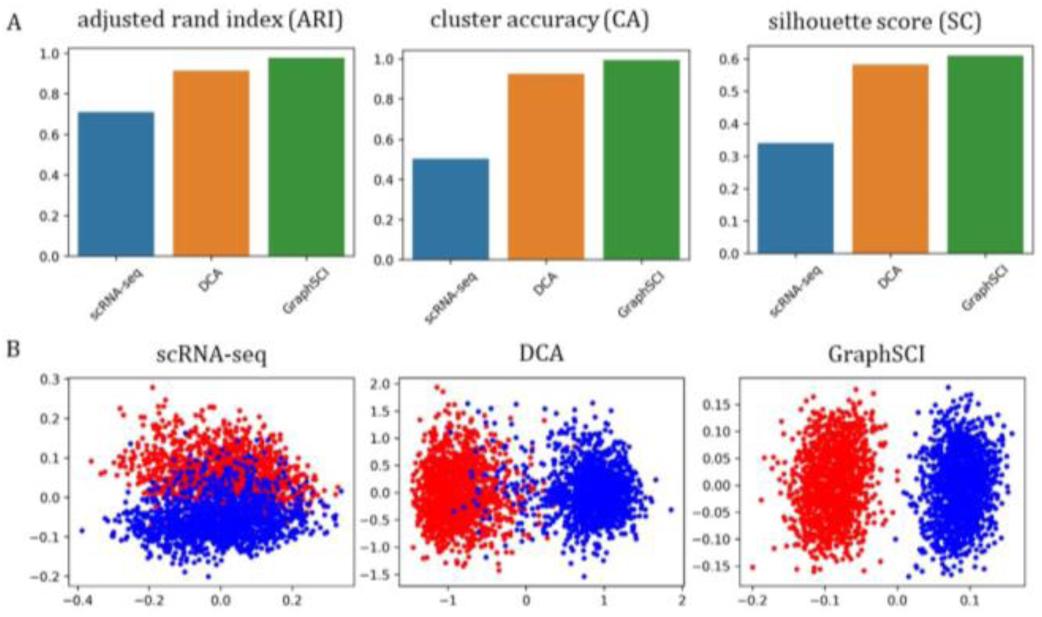
GraphSCI identifies cell types in simulated data with two cell groups. (A) The comparison of clustering performances of scRNA-seq, DCA and GraphSCI, measured by ARI, CA and SC. (B) The two principle components by PCA from simulated scRNA-seq data, imputed matrix by DCA and imputed matrix by GraphSCI. Each cell is colored by cell groups.

When tested on the SIM-T6 dataset with six cell groups, similar results can be obtained. As shown in Fig S1A, our method achieved 0.818, 0.859, and 0.034 for ARI, CA, and SC values, respectively. These are 20.5, 21.2, 30.8% higher than those by DCA with P-value of 0.023, and 106%, 131%, 78.9% higher than original data with P-value of 0.0027. The visualization consistently indicated that our method separates the six types of cells better than other methods (Fig S1B).

### GraphSCI recovers gene expression levels in bulk RNA-seq dataset

The efficacies of GraphSCI and DCA in recovering gene expression levels were further evaluated by a real RNA sequencing dataset. The RNA sequencing data was obtained from C. elegans development experiments by Eraslan et al^18^, which was used to simulate single cell RNA-seq data with dropout rates ranging from 50% to 70%. Since bulk RNA-seq data contains less noise than scRNA-seq, we used Pearson correlation coefficient (PCC) to evaluated the effectiveness of imputation on real RNA-seq dataset. As shown by Fig. 3, GraphSCI outperformed DCA in recovering the gene expression levels in real RNA-seq dataset. In details, the median of PCCs reached by GraphSCI in three datasets are 0.858, 0.809 and 0.782 respectively, comparing to the results 0.819, 0.759 and 0.750 achieved by DCA.

**Figure 3.**
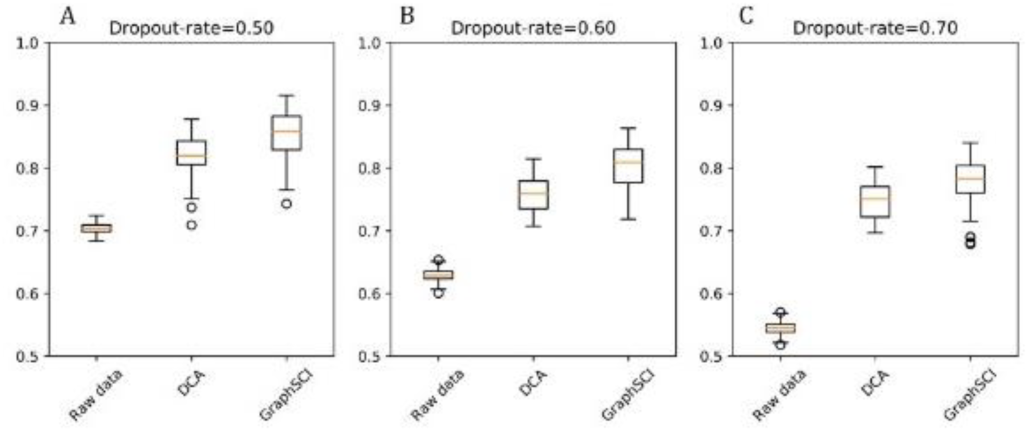
GraphSCI recovers gene expression levels in bulk RNA-seq data. Box diagram (A-C) depict the Pearson correlation coefficient between simulated data or imputed data and original data. And the box represents the interquartile range, the horizontal line in the box is the median, and the whiskers represent 1.5 times the interquartile range.

### GraphSCI recovers transcriptome dynamics in real single-cell data

Another key criterion to evaluate the imputation methods is their ability to recover transcriptome dynamics in real single cell dataset. Therefore, we applied our method to two real scRNA-seq datasets and compared it with others. The first dataset was obtained from mouse ES cells^19^, which were measured to analyze the heterogeneity of mouse embryonic stem cells in different stages after leukemia inhibitory factor (LIF) withdrawal. We selected four different LIF withdrawal intervals (0, 2, 4, and 7 days) and put all cells together as the input of imputation. The imputed data was clustered by t-SNE^20^. As shown by Fig. 4A, GraphSCI separated the four stages of mouse ES cells clearly except that a few blue samples were mixed with the yellow. In comparison, the clustering obtained from the scImpute and DCA method seriously mixed the blue samples with the yellow ones. As indicated by Figure 4B, ARI, CA, and SC of GraphSCI were significantly better than other approaches with P-values of 0.045, 0.031, respectively.

**Figure 4.**
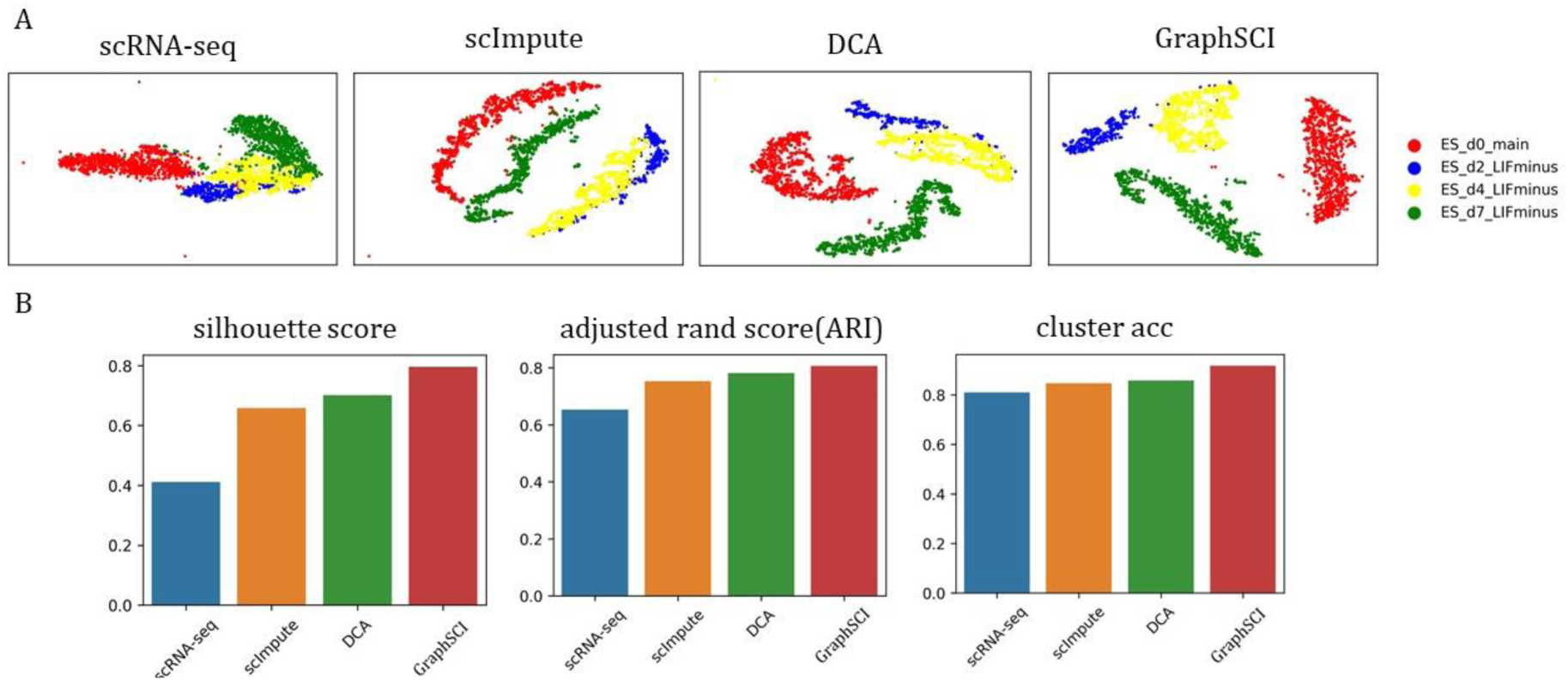
The performances on Mouse embryonic stem cells dataset. (A) shows the t-SNE visualization reproduced from scRNA-seq, scImpute, DCA and GraphSCI from left to right. (B) The comparison of clustering performances of scRNA-seq, scImpute, DCA and GraphSCI, measured by ARI, CA and SC.

GraphSCI and DCA^9^ were further applied to a large dataset generated by 10X scRNA-seq platform^21^, which is involved by the transcriptome of Peripheral blood mononuclear cells (PBMCs) from a healthy donor. The dataset contains 5247 Peripheral blood mononuclear cells of 11 cell types. Because the same type of cells has similar expression profiles, we selected 20% of PBMCs as an independent test set and trained the model on the remained data. The GraphSCI model was tested on the independent set and the imputed data was conducted with dimension reduction results by t-SNE^20^. From Figure 5A, we observed that the brown and dark samples were separated well in the low-dimension representation. The orange samples had a diving line with the other samples. In comparison, the results obtained by DCA were not observed with obvious differences with the black samples, red samples, and green samples. The results of ARI, CA, and SC on the independent test set also showed that GraphSCI outperformed other methods (Fig.5B). In details, our method achieved 0.472, 0.552, and 0.177 for ARI, CA, and SC values, respectively. These are 16.2, 9.9, 40.1% higher than those by DCA with P-value of 0.041.

**Figure 5.**
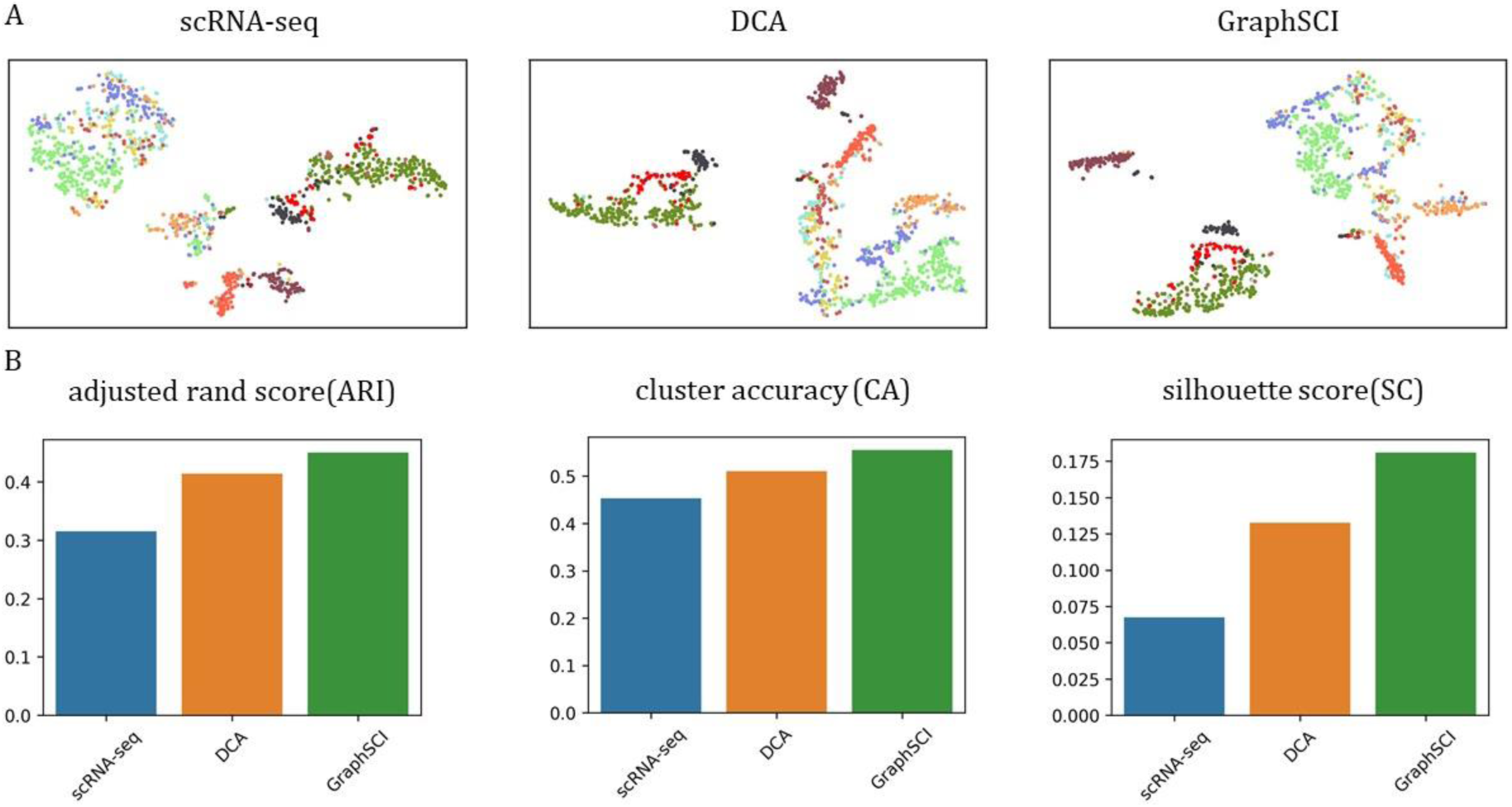
The performances on 5k peripheral blood mononuclear cells (PBMC) dataset. (A) shows the t-SNE visualization reproduced from scRNA-seq, DCA and GraphSCI from left to right. (B) The comparison of clustering performances of scRNA-seq, DCA and GraphSCI, measured by ARI, CA and SC.

### GraphSCI infers gene-to-gene relationships from scRNA-seq data

GraphSCI can not only impute gene expression data of scRNA-seq effectively, but also infer gene-to-gene relationships from the data. Due to the dropout events in raw scRNA-seq data^21, 22^, it is challenging to obtain accurate gene interactions directly from correlation coefficients between gene expression^23^. Here, we applied our method to raw scRNA-seq datasets and reconstructed gene relations during imputation. By compared with the known interactions from the STRINGdb^24^, the gene interactions constructed by GraphSCI had a precision of 0.713. Specifically, the true positive (TP) was 232647 and the false positive (FP) was 93812. Fig 6 shows the imputed gene-to-gene relationships obtained by Cytoscape^25^. Similar results were also observed in previous experiments on the mouse ES cells dataset. The constructed gene relations had a precision of 0.682 with 199291 true positives and 92924 false positives.

**Figure 6.**
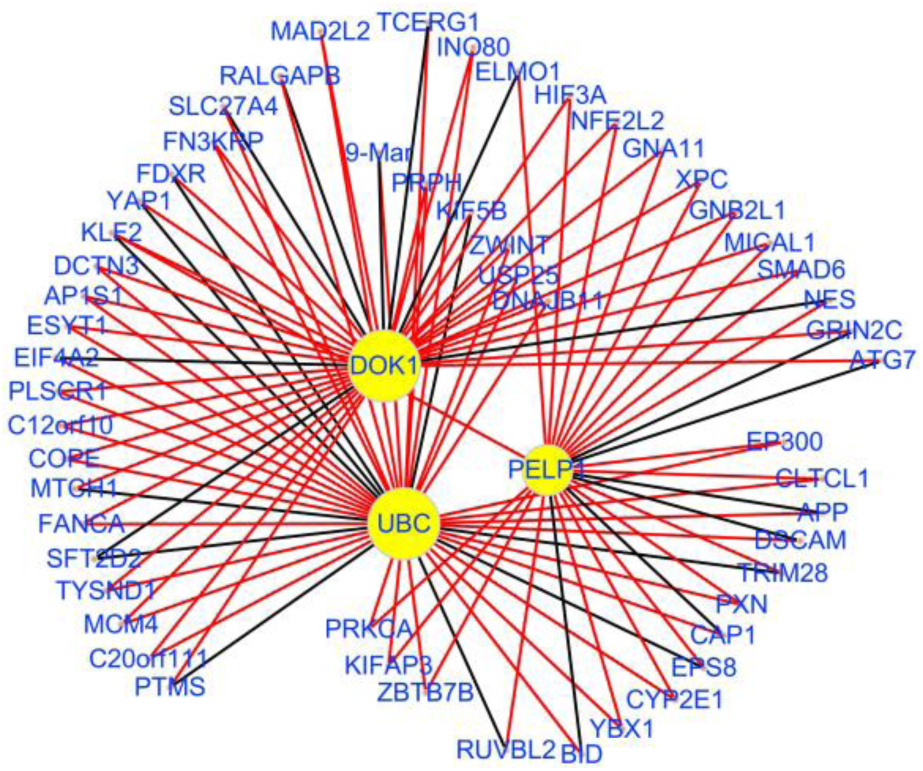
The gene-to-gene relationships after reconstruction. We selected the three genes with the highest degree and their common interactive genes. Compared to STRINGdb, the edges colored by red represent the correct gene-to-gene relationships we inferred and the black edges represent the false inferred relationships.

## Discussion

In this study, we presented a novel imputation method, GraphSCI, based on graph convolution networks, which are particularly suitable for single-cell RNA-seq data. Our method focused on imputing gene expression levels by integrating the gene expression with gene-to-gene relationships. By using gene-to-gene relationships as prior knowledge, this method avoided introducing excess biases during imputation and removed technical variations resulted from scRNA-seq.

GraphSCI was evaluated on simulated and real data, which was identified with the best performances on diverse downstream analyses in comparison with other methods. In simulated datasets, GraphSCI was found to outperform other methods on the data with small and large numbers of cells and cell types. The better performance of GraphSCI was further observed when it was applied in real dataset like bulk RNA-seq data and scRNA-seq data. In addition, another advantage of GraphSCI is its ability to infer the new gene-to-gene relationships, which is an absence of other methods.

To our best knowledge, our work is the first study to integrate gene-to-gene relationships into deep learning framework for imputation on scRNA-seq. It is also the first attempt to employ graph convolution network for learning the representation of gene-to-gene relationships in imputation study. Most importantly, extensive experiments were conducted on different kind of scRNA-seq datasets to demonstrate the superiority of our method.

Applications of graph convolution networks and exploiting gene-to-gene relationships for imputation, however, may also bring uncontrollable errors. For instance, the reliability of gene-to-gene relationships may influence the results of imputation. To solve this problem, we tried a variety of methods to build the gene-to-gene relationships, such as setting different thresholds to build edges or selecting original co-expressed samples to calculate Pearson correlation coefficients (PCC). We found that better performance could be achieved with the adjacency matrix obtained by selecting original co-expressed samples and PCC of >0.3 to determine edges.

Another challenge for real data is that the evaluation of imputation may be difficult because of the lacking of the ground truth. Therefore, we performed many clustering metrics, such as ARI, CA and SC, to describe the effectiveness and robustness of competing method, while we also utilized visualization to make the results clearer and more convincing.

The current GraphSCI was tested on datasets including simulated data and real data. The imputation power could be further improved with the increasing number of cells in the training set. Additionally, the deep learning networks by GraphSCI enables parallelization using GPUs to speed up training on large scRNA-seq datasets.

## Conclusions

In conclusion, our proposed method is shown to outperform competing methods over simulated and real datasets on diverse down-stream analyses. Furthermore, our study makes good use of the gene-to-gene relationships in the framework and infers new reliable relationships simultaneously. Altogether, we demonstrate that our proposed method is highly scalable and parallelizable via graphical processing units (GPU).

## Methods

The proposed model GraphSCI imputes gene expression levels in scRNA-seq data based on a combination of the graph convolution network and Autoencoder neural network, with the input of gene expression matrix *X* and gene-to-gene relationships *A*. Figure 1 shows the procedure of GraphSCI.

By using *M* single cells RNA-seq data with *N* genes, an undirected gene graph with gene expressions and gene-to-gene relationships can be constructed. Let 𝒩 and ℳ be a set of genes and samples respectively, an undirected gene graph can be denoted as *𝒢* = (𝒩, ℳ, *ℰ*), where *ℰ* is the set of gene-to-gene relationships. Thus, we introduce an adjacency matrix *A* ∈ ℝ^*N*×*N*^ and a gene expression matrix *X* ∈ ℝ^*N*×*M*^ for *𝒢*, with *A*_*ij*_ representing the edge of the *i*-th gene and the *j*-th gene and *X*_*ij*_ being the expression value with rows representing genes and columns representing cells. Table.1 summarizes our main notations for scRNA-seq data.

**Table 1.**
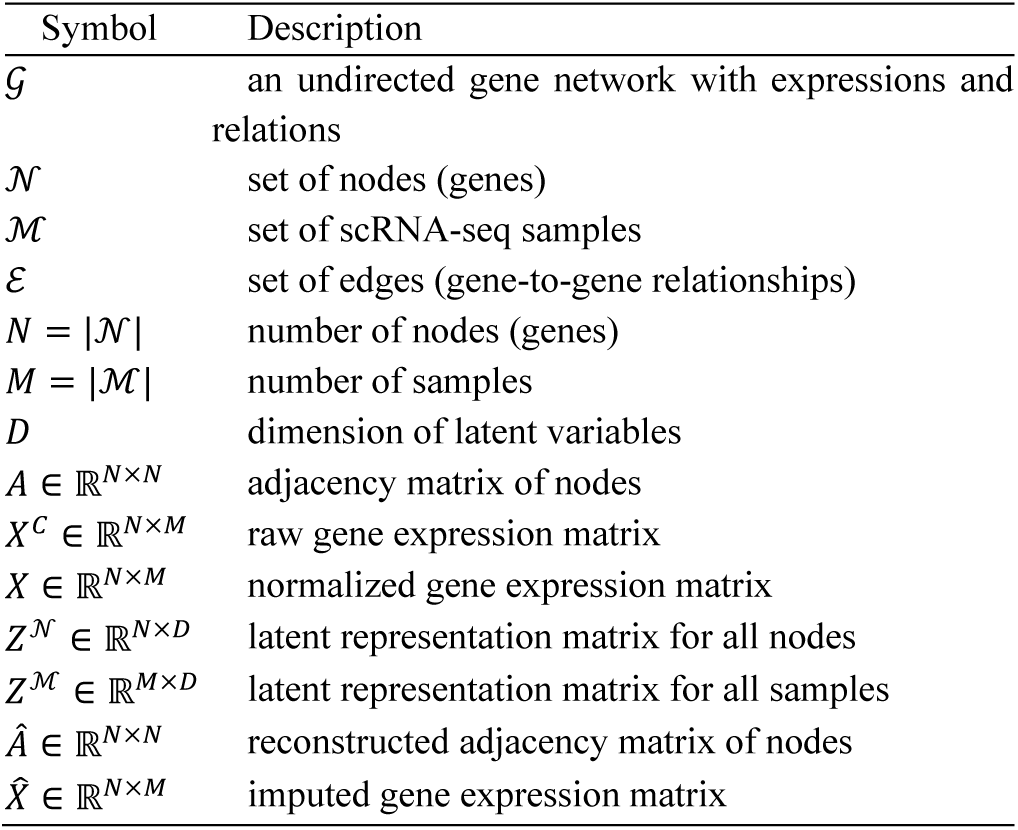
Main notations in our paper

### Data processing and normalization

There are two inputs to our proposed model: (1) a gene expression matrix *X* ∈ ℝ^*N*×*M*^, (2) an adjacency matrix *A* ∈ ℝ^*N*×*N*^, and our final goal is to construct an imputed gene expression matrix 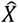 with the same dimensions. First, in raw scRNA-seq read count matrix *X*^*C*^, genes with no reads in any cell would be filtered out. Then, the library size of cell *i* is denoted as *l*_*i*_ and is calculated as the total number of read counts of cell *i*. The size factor *s*_*i*_ of cell *i* is *l*_*i*_ divide by the median of total counts per cell. Therefore, we make a normalized matrix *X* by taking the log transformation with a pseudo count and scale of the read counts:

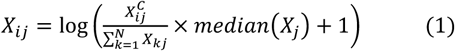

where *i* = 1,2, …, *N* representing each gene and *j* = 1,2, …, *M* representing each sample.

Secondly, we attempt to obtain the adjacency matrix *A* ∈ ℝ^*N*×*N*^ from a graph where genes are nodes and edges indicate genes which are likely to be co-expressed. For the simulated datasets generated from Splatter^16^, we introduce the adjacency matrix *A* ∈ ℝ^*N*×*N*^ by Pearson correlation coefficient (PCC) as:

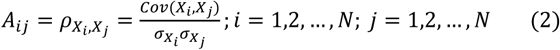

where *Cov*(*X, Y*) and *σ*_*X*_ is the covariance between *X* and *Y* and the standard deviation of *X* respectively.

Moreover, the adjacency matrix *A* ∈ ℝ^*N*×*N*^ from real datasets can be created by Pearson correlation coefficient as mentioned above and protein-protein interaction (PPI) databases, like STRINGdb^24^.

### Imputation based on graph convolution network

The pre-processed gene expression matrix and adjacency matrix are treated as the input for GraphSCI. Two neural network models, i.e., the inference model *f*_*ϕ*_ and the generative model *g*_*φ*_ were used to constructed the model for the probabilistic encoder *q*_*ϕ*_ and probabilistic decoder *p*_*φ*_ respectively, to preform gradient descent for learning all trainable parameters.

To infer the embeddings of cells and genes, we apply a two-layer graph convolution network and a two-layer fully connected neural network mapping the adjacency matrix A and the gene expression matrix X to the low-dimensional representations of the posterior distribution (i.e. Gaussian distributions and ZINB distributions) respectively. In particular, the two-layer GCN is defined as:

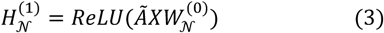

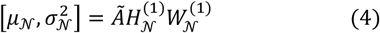

where *μ*_𝒩_ and 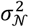 are the mean and variances of the learned Gaussian distribution parameters, *ReLU*(·) = max (0, ·) is the non-linear activation function, 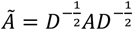 is the symmetrically normalized adjacency matrix with the *𝒢*′ s degree matrix *D* _*ii*_ = ∑ _*j*_ *A* _*ij*_, and 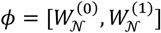 are the trainable parameters of GCN layers.

The two-layer fully connected layers for inferring ZINB distribution of single cell samples are defined as:

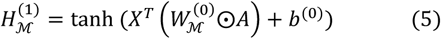

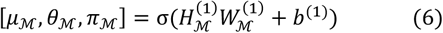

where *μ*_ℳ_, *θ*_ℳ_ and *π*_ℳ_ are the parameters of the ZINB distribution: mean, dispersion and dropout probability, the operation ⊙ is the Hadamard (element-wise) product, tanh(·) and s(·) are the activation functions and 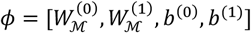 are the trainable parameters of fully connected layers.

In particularly, the ZINB distribution is applied for count data that exhibit over-dispersion and excess zeros, which is parameterized with the mean (*μ*) and dispersion (*θ*) of the negative binomial distribution as well as the dropout probability (*π*) representing the probability of zeros (dropout events). A count matrix X that is ZINB-distributed with (*μ, θ, π*) is denoted:

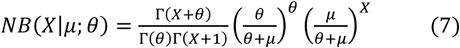

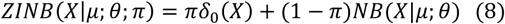

where Γ(x) and *δ*_0_(*x*) is the Gamma function and Dirac function respectively. Therefore, we could estimate the parameters *μ, θ, π* of ZINB distribution from the hidden layer in Eq. (6):

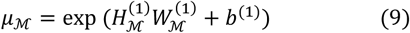

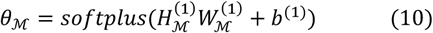

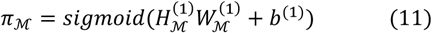

where exp(·) is the exponential function and softplus(·) and sigmoid(·) are the non-linear activation functions.

After having obtained the parameters of the learned distributions, the reparameterization method could help us transform the latent variables 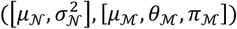 to deterministic variables, denoted as *Z*^𝒩^, *Z*^ℳ^. Therefore, the generative model in our framework could decode from the deterministic variables *Z*^𝒩^ and *Z*^ℳ^ to generative random variables, where the gene expressions and gene-to-gene relationships can be reconstructed.

Specifically, given embeddings of gene *i* and cells *j*, we compute 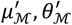 and 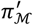 by:

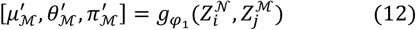

where 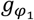 is a neural network for reconstructing gene expression matrix and *φ*_1_ is the trainable parameter in 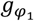. Then an imputed gene expression 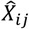 can be generated by the following process:

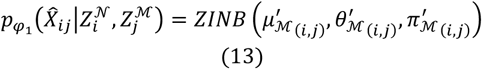

where 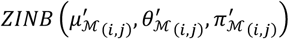 is the ZINB distribution parameterized by 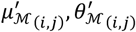 and 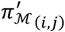 and 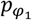 is the probabilistic decoder given the latent embeddings 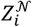 and 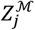.

Therefore, we could implement the generative model 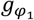 by:

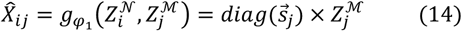

where diag(·) is the diagonal matrix constructed by the vector (·) and 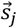 is the size factor of cell *j*.

Similarly, given embeddings of two genes *i* and *j*, we can compute 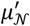 and 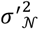 by:

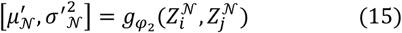

where 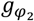 is a neural network for reconstructing gene-to-gene relationships and *φ*_2_ is the trainable parameter in 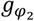. Then an observed edge between two genes *i* and *j* can be generated by:

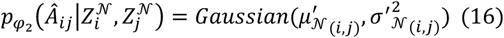

where 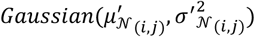 is the Gaussian distribution parameterized by 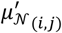 and 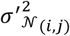 and 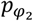 is the probabilistic decoder given the latent embeddings 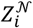 and 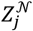.

The generative model 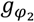 to reconstruct gene-to-gene relationships could be defined as:

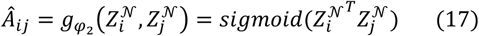

where sigmoid(·) is the sigmoid function.

### Optimization

The optimization was performed to obtain accurate embeddings of both genes and cells in an unsupervised way. For this purpose, *Z*^𝒩^ and *Z*^ℳ^ were optimized by the variational lower bound *ℒ*:

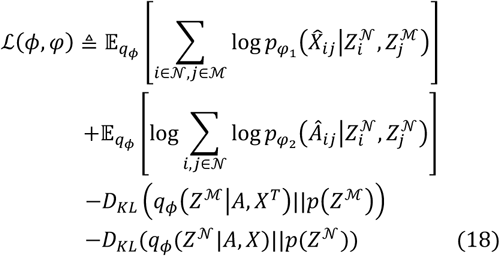

where 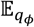 is the cross entropy function with the probabilistic distribution *q*_*ϕ*_ and *p*_*φ*_ and 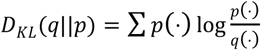 is the Kullback-Leibler (KL) divergence between q(·) and p(·). In the above equation, *q*_*ϕ*_(*Z*^ℳ^|*A, X*^*T*^) and *q*_*ϕ*_(*Z*^𝒩^|*A, X*) is defined as the probabilistic encoder with the input of *A, X*^*T*^ and *A, X* respectively, aiming at producing the representations *Z*^𝒩^, *Z*^ℳ^. Similarly, 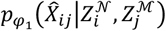 and 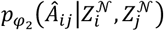 are the probabilistic decoders for construct the imputed gene expression matrix 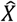 and gene-to-gene relationships *Â*. Furthermore, the KL divergence in optimization could be interpreted as the regularization to make the predicted posterior distributions closer to the prior distributions *p*(*Z*^ℳ^), *p*(*Z*^𝒩^).

With the help of reparameterization trick, we could represent the distributions with deterministic variables:

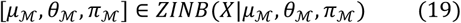

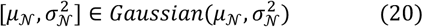

These deterministic variables are differentiable and capable to be calculated in backpropagation process. We could directly derivate Eq. (18) based on Monte Carlo estimates:

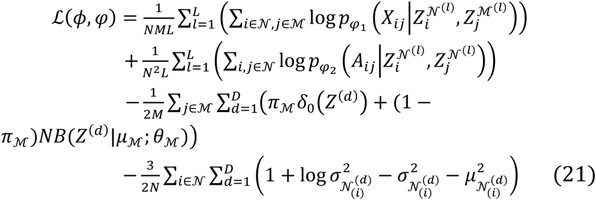

Therefore, with the optimization, the gradient-based optimization techniques can be used to train the end-to-end model.

### Evaluation metrics

Clustering performance can be measured by the clustering metrics: adjusted Rand index^26^ (ARI), clustering accuracy (CA) and Silhouette Coefficient^27^ (SC).

The adjusted Rand index (ARI) is the corrected-for-chance version of the Rand index. The Rand index is a measure of the similarity between two data clustering and the ARI is adjusted for the chance grouping of elements. Given a set of n samples, the two clusters of these samples are *V* = {*V*_1_, *V*_2_, …, *V*_*r*_} and *U* ={*U*_1_, *U*_2_, …, *U*_*t*_} and *n*_*ij*_ is defined as *n*_*ij*_ = |*V*_*i*_ ∩ *U*_*j*_|. Let 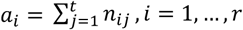 and 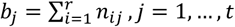, the ARI could be defined as

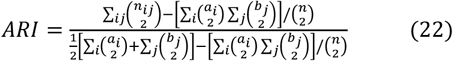

The CA is defined as the accuracy of the clustering assignments. Given a sample *i*, let *s*_*i*_ be the ground truth label and *r*_*i*_ be the assignments of clustering, then the CA is

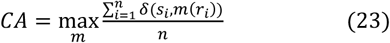

where *n* is the number of samples, *m* is the set of one-by-one mapping between clustering assignments and true labels and *δ*(x, y) = 1 if x = y otherwise 0.

The SC measured the similarity between a single cell and its cluster. The silhouette ranges from −1 to +1, where a high value indicates that the object is well matched to its own cluster. It could be defined as

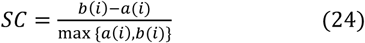

where *a*(*i*) is the mean distance between sample *i* and all other samples in the same cluster and *b*(*i*) is the minimum distance of sample *i* to all points in any other cluster.

### Simulated datasets

Our simulated data are generated by Splatter^16^ R package, a widely used package for simulating the scRNA-seq count data. First, we simulated a dataset with two cell groups, 2000 cells of 3000 genes by setting 27% of data values to zero mimicking dropout events. During the simulation, we set the parameter *dropout. shape* = −1, *dropout. mid* = 0 and *de. fracScale* = 0.3 for simulating the dropout events and the other parameters are set to default values. Hence, we could obtain the true counts before dropout and the raw counts after dropout, which are the simulated scRNA-seq data. Furthermore, we simulated a complex dataset of 3000 cells by 5000 genes to evaluate the robustness of our model, The 3000 cells are divided into six groups and the parameter were set to *dropout. shape* = −1, *dropout. mid* = 0, *de. fracScale* = 0.3 and the other parameters with default values.

### C. elegans time course experimental data

We obtain the bulk transcriptomics data from the supplementary material of Franceconi. et al, which contains 15855 detected genes during 12 hours of C. elegans development^18^. We analyzed the dataset after simulating single-cell transcriptomics dropout noises and the bulk transcriptomics data can be the ground truth for evaluation. Hence, we compared our method with the existing method DCA by Pearson correlation coefficient.

### Mouse embryonic stem cells data

Klein. et al. profiled the single-cell transcriptomics by droplet-microfluidic approach and applied it on embryonic stem cells^19^. They analyzed the heterogeneity of mouse embryonic stem cells differentiation after leukemia inhibitory factor (LIF) withdrawal. Here, we selected the four different LIF withdrawal intervals (0, 2, 4, 7 days) and construct a scRNA-seq dataset with 2717 cells of 24175 detected genes. And the cell types are determined by the intervals of LIF withdrawal.

### 5k peripheral blood mononuclear cells (PBMC) from a healthy donor

The dataset was provided by 10X scRNA-seq platform^21^, profiling the transcriptome of the peripheral blood mononuclear cells (PBMCs) from a healthy donor. The total number of cells was 5247 after filtering process and the cell types were identified by graph-based clustering on the platform.

### Human Embryos cells scRNA-seq data

Xue et al. performed a comprehensive analysis of transcriptome dynamics by weighted gene co-expression network analysis^28^. Therefore, we could obtain the dataset containing 30 samples from oocyte to morula in human embryos samples from their experiments. Here, we utilized the dataset to demonstrate the effectiveness of our method on inferring the gene-to-gene relationships.

### Implementation

We implemented the proposed model with Tensorflow 1.11.0^29^. In the training process, we utilized the Adam^30^ optimizer with an initial learning rate of 0.01 and allowed it to decay exponentially with decay_rate = 0.9 and decay_steps = 50 during learning. The total loss and learning rate decreased with epoch during training as shown in supplementary Fig. 2. The hidden layers of encoders were set as 16 neurons and we use a 32-dimensional of embedding latent variables in all experiments, denoted as *D*. To alleviate overfitting, we implemented the regularization methods such as dropout and L2 regularization. Dropout^31^ rate 0.2 was applied on the inference model and the coefficient of L2 regularization was 0.001. We explored hyper-parameters in a wide range and find the above hyper-parameters yields the highest performance, as supplementary Fig. 3 shown. All experiments are repeated for 5 times, each with a different random seed. The implementation is publicly available at https://github.com/GraphSCI/GraphSCI.

## Data availability

The scRNA-seq datasets used in this manuscript are publicly available and their details are summarized in Table 2. The C. elegans time course experimental data was provided by the supplementary material of Francesconi. et al. The mouse embryonic stem cells data was downloaded from GSE65525. The 5k peripheral blood mononuclear cells (PBMC) from a healthy donor was provided by the 10X scRNA-seq platform and the website of the data is https://support.10xgenomics.com/single-cell-gene-expression/datasets/3.1.0/5k_pbmc_protein_v3. The human embryos cells scRNA-seq data was downloaded from GSE44183.

**Table 2.** The summarization of datasets in this manuscript

| Datasets | Sample size /<br>cell number | Number of<br>genes | Number of cell<br>types |
| --- | --- | --- | --- |
| SIM-T2 | 2000 | 3000 | 2 |
| SIM-T6 | 3000 | 5000 | 6 |
| C. elegans time-course | 206 | 15855 | - |
| Mouse ES cells | 2717 | 24175 | 4 |
| 5K PBMC | 5247 | 33570 | 11 |
| Human ES cells | 30 | 14766 | - |

## Acknowledgements

This work has been supported by the National Key R&D Program of China (2018YFC0910500), National Natural Science Foundation of China (U1611261, 61772566, and 81801132), Guangdong Frontier & Key Tech Innovation Program (2018B010109006, 2019B020228001) and Introducing Innovative and Entrepreneurial Teams (2016ZT06D211).

## Author Contributions

J.R., X.Z. and Y.Y. contributed concept and implementation. J.R. designed experiments. J.R. was responsible for programming. All of the authors contributed to the interpretation of results. J.R., H.Z. and Y.Y. wrote the manuscript. All of the authors reviewed and approved the final manuscript.

